# Brain heating induced by near infrared lasers during multi-photon microscopy

**DOI:** 10.1101/057364

**Authors:** Kaspar Podgorski, Gayathri Ranganathan

**Affiliations:** Janelia Research Campus, Howard Hughes Medical Institute, VA, USA 20147

**Author notes:** Contact, 1-(. **New and Noteworthy** New optical tools are transforming neuroscience, allowing increasingly comprehensive control and readout of neural circuits. However, emerging parallel microscopy methods rely on higher total illumination powers than conventional techniques, raising concerns about heating and its effects on the brain. This study characterizes these effects under serial and parallel illumination schemes, allowing researchers to predict power limits for their experiments. Even under serial illumination, usable power can be limited by heating rather than nonlinear damage.

**Keywords:** Multifocal, multiphoton, hyperthermia, nanothermometry

## Abstract

Two-photon imaging and optogenetic stimulation rely on high illumination powers, particularly for state-of-the-art applications that target deeper structures, achieve faster measurements, or probe larger brain areas. However, little information is available on heating and resulting damage induced by high-power illumination in the brain. Here we used thermocouple probes and quantum dot nanothermometers to measure temperature changes induced by two-photon microscopy in the neocortex of awake and anaesthetized mice. We characterized heating as a function of wavelength, exposure time, and distance from the center of illumination. Although total power is highest near the surface of the brain, heating was most severe hundreds of microns below the focal plane, due to heat dissipation through the cranial window. Continuous illumination of a 1mm^2^ area produced a peak temperature increase of approximately 1.8°C/100mW. Continuous illumination with powers above 250 mW induced lasting damage, detected with immunohistochemistry against Iba1, GFAP, heat shock proteins, and activated Caspase-3. Higher powers were usable in experiments with limited duty ratios, suggesting an approach to mitigate damage in high-power microscopy experiments.

## Introduction

Modern neuroscience has been transformed by tools that use light to observe and manipulate brain cells (Chen et al., 2013; Hochbaum et al., 2014; Emiliani et al., 2015). Optical methods allow spatially dense readout and optogenetic control of neurons *in vivo* with minimal perturbations of the brain. Efforts are underway to optically address larger numbers of cells more rapidly, for example, to more comprehensively record and manipulate rich patterns of activity in 3-dimensional space that transmit information in the brain (Emiliani et al., 2015).

Addressing optical effectors and reporters at high resolution within intact tissue usually relies on multiphoton absorption (e.g. Packer et al., 2015; Papagiakoumou et al., 2013; Voigt et al., 2015; Bahlmann et al., 2007; Horton et al., 2013). The throughput of these methods increases with higher laser power, by addressing more foci in parallel or increasing signal rates from each. In two-photon imaging, doubling the laser power quadruples the signal rate achievable from a given focal pattern. However, light absorption heats brain tissue, which can change neural function (Hodgkin and Katz, 1949; Milburn et al., 1994; Aronov et al., 2011; Stujenske et al., 2015; Kiyatkin, 2007). Heating is of particular concern for multiphoton techniques, because simultaneous absorption of two coherent photons occurs with low probability and requires high light intensities. To reduce average power, brief and high-intensity laser pulses are used. Pulsed illumination increases two-photon absorption efficiency due to its square dependence on intensity, but also increases photodamage and bleaching rates, which depend on intensity raised to powers greater than 2 (Drobizhev et al., 2014; Kalies et al., 2011; Patterson and Piston, 2000). This nonlinear photodamage limits the peak power that can be used in living tissues, creating a practical lower bound on the average power (and therefore heating) needed to achieve a given signal rate.

Heating can perturb brain activity in several ways. Virtually all biophysical properties are temperature-dependent, including the waveform of the action potential (Hodgkin and Katz, 1949) and channel conductances (Milburn et al., 1994; Shibasaki et al., 2007; Wells et al., 2007). Illumination can depolarize cells directly through multiple mechanisms (Fork, 1971; Hirase et al., 2002; Shapiro et al., 2012). Even relatively low illumination powers can increase neuronal firing rates (5mW at 532nm; Stujenske et al., 2015). Neural circuit dynamics are also affected by both manipulations and natural variations in brain temperature (Aronov and Fee, 2012; Aronov et al., 2011), and febrile seizures occur in children with elevated brain temperatures (Pavlidou et al., 2013). Heating can cause lasting damage to tissues, and the brain is especially sensitive to such damage (Dewhirst et al., 2003). Extreme heating induces edema, denaturing and coagulation of proteins, inflammatory responses, and cell death (Chen et al., 2000; Dewhirst et al., 2003; Lepock, 2003). Onset of damage occurs at temperatures ranging from 40-44°C depending on duration and conditions of exposure, and different brain tissues are differentially sensitive (Chen et al., 2000; Kiyatkin, 2007). Hyperthermia also aggravates other neurological insults (Kim et al., 1996), while hypothermia is neuroprotective (Barone et al., 1997; Yenari and Han, 2012).

Despite years of applications, measurements of brain heating and associated damage induced during two-photon excitation are not available. Such measurements are needed because neuroscientists continue to advance imaging and photostimulation methods to address larger numbers of neurons in parallel, at higher speeds, and deeper in tissue. Here we used thermocouple measurements and quantum dot spectrometry to characterize heating under two-photon microscopy conditions, and ascertain practical limits to average power used for these experiments.

## Methods

All experimental protocols were conducted according to U.S. National Institutes of Health guidelines for animal research and approved by the Institutional Animal Care and Use Committee at Janelia Research Campus (protocol 14-115).

**Mice.** Experiments were performed with C57Bl6J mice (Charles River Laboratories). One experiment (Fig. 6) involved GP4.12 mice (Dana et al., 2014). Both male and female mice, between 8 and 22 weeks of age, were included in studies. All surgery was performed under isoflurane anaesthesia (1.25-2%v/v). Anaesthetized mice were heated to maintain a body temperature of 37°C during all procedures.

**Thermocouple Probes.** Flexible thermocouple probes (IT24P, Physitemp) were mounted in rigid capillary glass, leaving 2mm of the probe exposed at the tip. The maximum diameter of the probe entering the brain was 220 |im. Mouse brain and body temperature were recorded simultaneously with a two-channel thermometer/calibrator (CL3515R, Omega).

**Thermometry in freely moving mice.** A 4mm incision was made in the skin above visual cortex. Through this incision, a hole was drilled above visual cortex (2.2mm lateral and 1mm anterior of lambda) with a #4 burr. The thermocouple was inserted to a depth of 100 |im below the dura surface using a micromanipulator, and the skull was sealed with a small quantity of dental cement. The incision was closed around the thermocouple wire with veterinary adhesive (VetBond). Brain temperature was monitored for 30 minutes after recovery from anaesthesia as mice moved freely in their home cage. Temperatures at 30 minutes are reported.

**Cranial Windows.** Partial windows used with thermocouple measurements consisted of a single 3.5mm diameter circular #1 coverslip, cut along one edge. A 3mm diameter craniotomy was made over visual cortex and the surrounding bone was thinned. A drop of low-temperature gelling agarose (1%in cortex buffer) was placed on the brain surface and the window pressed over the craniotomy with light pressure. The window was cemented in place with dental acrylic and a headbar with a circular well to hold immersion fluid was cemented to the skull surrounding the window. For lateral distance measurements (Fig. 1h–i), a craniotomy spanning 7mm anterior-posterior adjacent to the midline was used, with larger window glass. A small hole was made in the dura near the edge of the window with a patch pipette to allow the thermocouple to penetrate. A single penetration was used per mouse. The thermocouple was inserted into the brain as close to the edge of the glass window as possible, and advanced below the cranial window to the appropriate depth. Awake experiments were performed one day after window implantation. To prevent agarose from drying during recovery from surgery, the craniotomy was sealed with Kwik-Cast silicone elastomer (World Precision Instruments), which was removed before imaging. Room temperature saline was used as immersion fluid and allowed a minimum of 5 minutes to reach equilibrium temperature before thermometry.

**Figure 1.**
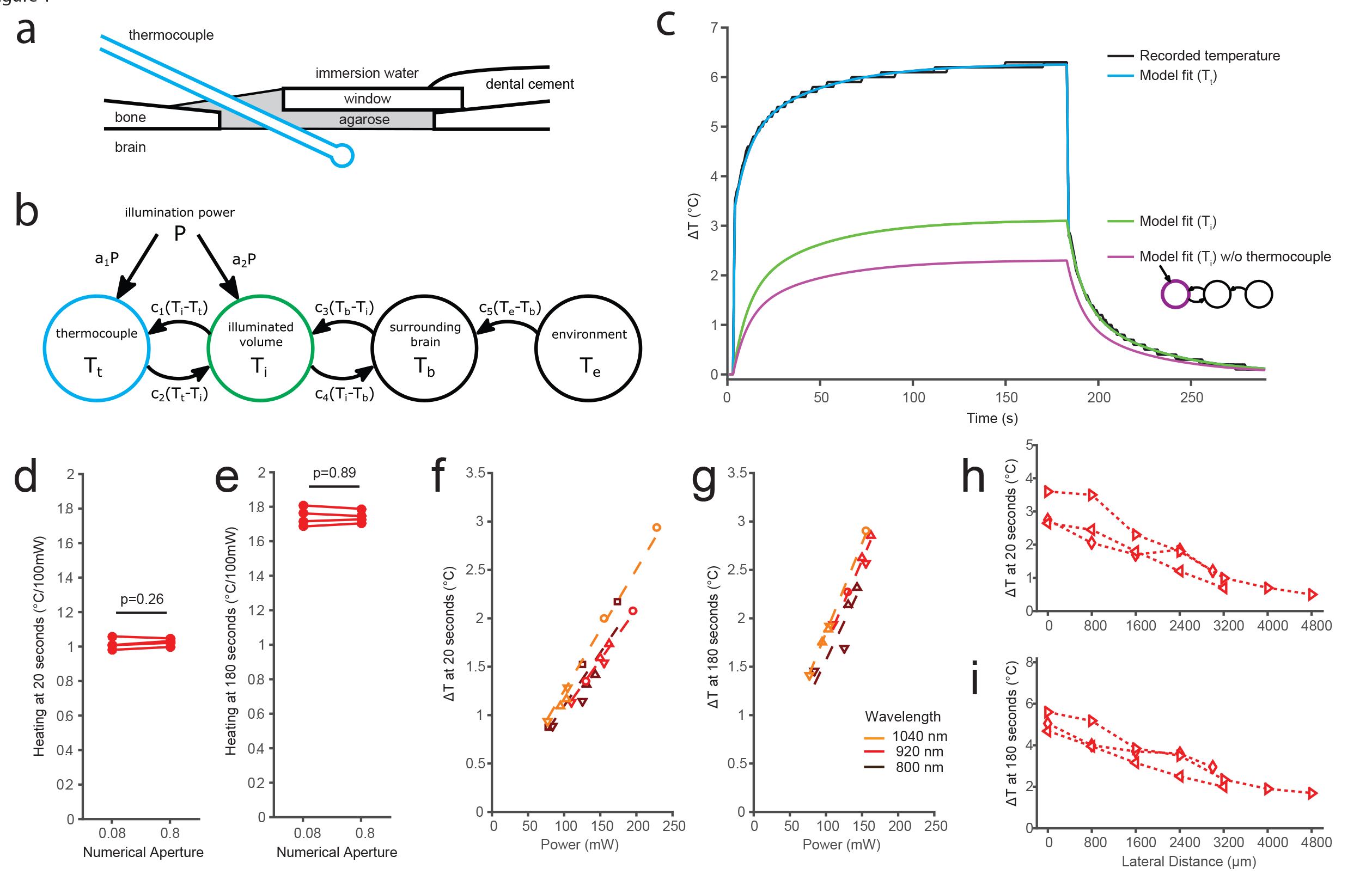
Thermocouple measurements of laser-induced brain heating. a) Experimental schematic. A thermocouple (220 μm diameter) was inserted under a partial cranial window into the brain region being imaged. Not to scale. b) The 4-compartment ODE used to model temperature changes near the thermocouple. c) Measured heating (2 replicates, black) at 500 μm depth with 920 nm, 155mW illumination. Corresponding model fits for the thermocouple (blue), illuminated brain volume (green), and illuminated brain volume in the absence of the thermocouple (purple). Inset, schematic of the ODE model in the absence of the thermocouple. (d,e) Heating coefficients (slope of lines in f-g) measured for illumination with highly focused (NA 0.8) or weakly focused (NA 0.08) light at powers of 0-230 mW (Paired t-tests). f,g) Measured heating at 500 μm depth after 20 (f) or 180 (g) seconds of illumination at 500 μm depth. (h,i) Measured heating at lateral distances from the center of the scanned area, with 920nm, 350 mW illumination. Symbol shapes in f-i denote biological replicates; each symbol denotes the same mouse throughout the figures. N=4 mice (d,e,f), N=3 mice (g,h,i).

Cranial windows used in QDot spectrometry and histology experiments were made from two layers of glass (3.5 mm and 4.5 mm diameter) bonded to each other. A 3.5mm diameter craniotomy was made over visual cortex and surrounding bone was thinned. The window was implanted such that the edge of the larger window rested on the bone surface and smaller window rested inside the craniotomy (Fig. 2b).

**Figure 2.**
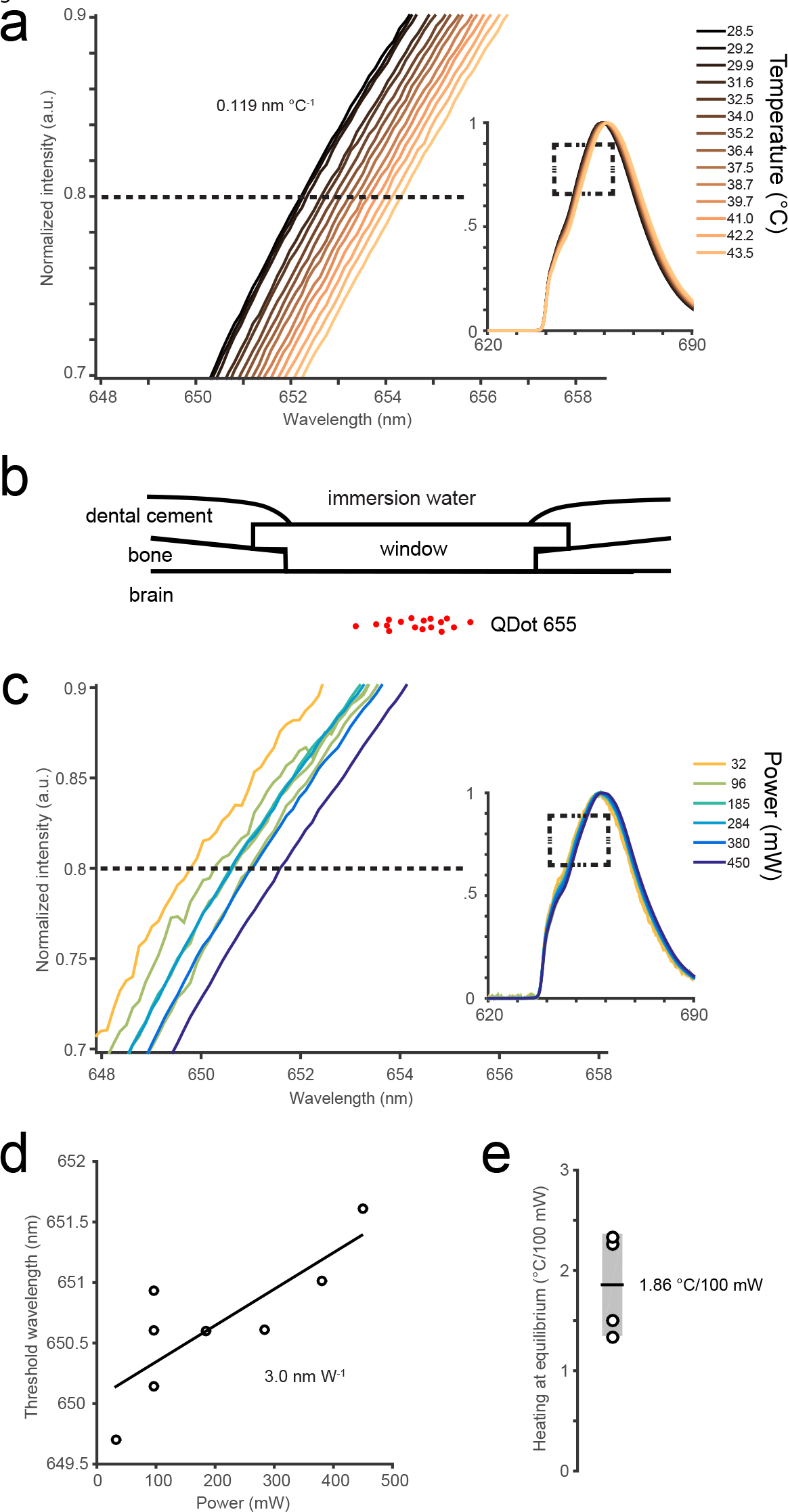
Quantum dot measurements of laser-induced brain heating. a) QDot 655 two-photon excited fluorescence emission spectra measured *in vitro*. Plots show the magnified spectrum in the vicinity of the intensity threshold (80%of maximum) used to identify spectral shifts. The inset shows full spectra. Spectra are red-shifted with increasing solution temperature. b) Schematic for *in vivo* temperature measurements. Qdot 655 nanocrystals were nanoinjected 250 μm below the dura, and a cranial window implanted above. Not to scale. c) Example two-photon excited emission spectra measured *in vivo* at different illumination powers. The inset shows full spectra. d) Threshold wavelengths at different illumination powers for the experiment in (c). e) Measured dependence of steady state temperature on illumination power for all experiments (N = 4 mice). Shaded region denotes 95%confidence interval for the group mean.

**QDot Thermometry.** Qdot 655 ITK Amino PEG-coated quantum dots (Invitrogen) were nanoinjected (40nL, 1 μM) into anterior visual cortex. A cranial window was implanted 1 week after injection, and spectrometry was performed 1 week after window implant. The microscope (920 nm excitation) was focused 250um below the dura, and detected light was passed through an emission filter (HQ675-70m-2P, Chroma) and coupled to the optical fiber input of a spectrometer (Acton SP2300i, Princeton Instruments). Laser scanning started at least 3 minutes before spectrum acquisition onset to ensure the temperature in the focus reached steady state. Control spectra to determine spectral shifts with temperature (shift: 0.119 nm °C^−1^, r^2^=0.998) and illumination power (shift: 1.41 nm W^−1^, r^2^ = 0.983) were obtained in a sealed glass well (~50 μL) immersed in water. Qdot 655 solution (750 nM) was pumped through the well with a syringe (1mL/30s) while the emission spectrum was measured. Temperature within the well was monitored with an embedded thermocouple. Spectra were acquired with exposure times of 30 (*in vitroo*) and 120-1000 s (*in vivo*). For temperature calibrations, the dye temperature ranged between 28.5-44.8°C. Illumination power calibrations were obtained at 28.3°C.

**Two-photon imaging and heating.** Experiments were performed using three two-photon microscopes with similar designs, incorporating resonant (~8kHz) galvanometer scanners, and controlled by Scanimage software (Vidrio Technologies). 512x512 images (1mm^2^) were acquired at 30±0.1 Hz using bidirectional scanning. The time duty cycle was 70%, with the beam blanked for 15%of a cycle at each extremum of the sinusoidal fast axis scan. This corresponded to an average pixel dwell time of ~90ns per 1.95um pixel. The proportion of time that the beam is blanked affects the spatial uniformity of the scan and the peak power at the focus; a longer beam blanking period implies a higher illumination intensity for a shorter period of time at each sample point. Chameleon (Coherent) lasers were used for studies in Figs. 2 and 4, and an Insight DS+ (Spectraphysics) laser for studies in Figs. 1,3, and 4. The three lasers used had similar repetition rates of 80MHz and pulsewidths of approximately 120 fs. This corresponds to a peak power of ~10kW at an average power of 100mW. We performed experiments with both highly focused and weakly focused beams, with numerical aperture (NA) 0.8 and 0.08 respectively. A weak focus was achieved by inserting a lens into the excitation path prior to the scan mirrors, decreasing the beam diameter at the objective pupil (‘underfilling the objective’). The inserted lens (f=250mm or 400mm, depending on the microscope) was chosen to create a focus 25 mm beyond the scan mirrors. This resulted in a modeled focal shift of 170um, a scan demagnification of 2%, a focal spot Airy diameter of 14 μm at 920nm and a decrease in peak intensity at the focus of approximately 100x compared to the fully filled back aperture. The decrease in two-photon excitation and increase in spot size were confirmed by imaging Qdot-injected mice with and without the lens in place (FWHM of putative subresolution emitters: 6.3±0.1 μm). We assessed the effect of NA on heating by alternating trials with or without the defocus lens in place, while adjusting the focal depth and scan area to compensate for the changes induced by the lens. In all experiments, the focal plane of the microscope was aligned to 250 μm below the dura, which was detected by autofluorescence. Laser powers at each wavelength and power setting were measured after the objective with an optical power meter (FieldMate; Coherent) before each experiment. Scanimage was used to scan the laser over a 1×1 mm square field while recording thermocouple temperature. The thermocouple was placed at either 150 μm or 500 μm below the dura, and the field of view was centered on the thermocouple or, for Fig. 1h–i, displaced laterally by the indicated distance. Wavelengths of 800, 920, and 1040 nm were used.

**Modeling.** A 4-compartment model was used to fit recorded thermocouple heating data. Temperatures in the model evolve according to a set of ordinary differential equations, the couplings in Fig. 1b, with illumination *P* as a time-varying input. The model has 7 parameters (*a_1_-a_2_,c_1_-c_5_*). The initial temperature of all compartments was set to the measured thermocouple temperature before heating onset (Δ*T*= 0). Five free parameters (*a_1_-a_2_,c_3_-c_5_*) were fit with the Matlab system identification toolbox function idgrey. Parameters *c_1_* and *c_2_* were determined by the following experiments: *c_1_*, the rate of temperature transfer from surrounding tissue to the thermocouple, was measured by rapidly inserting the thermocouple into brain tissue while recording thermocouple voltage at 10kHz. Traces recorded in this way were fit with a modified ODE system without laser heating to determine *c_1_*. *c_2_*, the rate of temperature transfer from the thermocouple to surrounding tissue, was fixed by heating the thermocouple directly and measuring resulting changes in surrounding tissue temperature after heat offset, again by fitting a modified ODE model without laser heating.

A finite differencing model was adapted from Stujenske et al. (2015) to simulate heating by a scanned objective focus, heat conduction, metabolic heating, and cooling by blood perfusion. The model was modified 1) to focus incoming light rays to a point that was then scanned in space, 2) to simulate a heterogeneous sample including the 350 μm glass window and immersion water, and 3) to improve numerical stability. The boundary conditions of the model were altered to fix the temperature of the immersion fluid at 25°C 1.5 mm above the glass surface. Parameter settings: Voxel size 0.01 mm (light diffusion) 0.03mm (heat diffusion), time step 0.16 ms, wavelength 920nm. Material constants for glass and water were obtained from published tables. Modifications to the code are available from KP.

**Immunohistochemistry.** Histological experiments were performed 3 weeks after window implantation. Awake headfixed mice were exposed to one of two heating protocols: 1) continuous scanning for 20 minutes, or 2) 10 second scans at 30 second intervals for 1 hour. Both protocols resulted in the same total amount of illumination. After 16 hours, mice were transcardially perfused with 4%paraformaldehyde in 0.1 M phosphate buffer (pH 7.4). Brains were postfixed in the same solution for 4 hours, followed by wash in phosphate buffered saline (PBS, pH 7.4). Coronal sections were cut through the center of the heated region and incubated with primary antibodies, supplemented with 2%bovine serum albumin and 0.4%Triton X-100. Alternating slices were labeled for Iba1 and GFAP, or HSP and Caspase-3. Slices were washed three times and incubated with species-appropriate secondary antibodies (1:500 dilution) conjugated to Alexa Fluor 488 (Iba1,HSP), Alexa Fluor 594 (GFAP), Cy3(Caspase-3), or Cy5 (Iba1, HSP in Figure 6b) washed again, and mounted for imaging. Primary antibodies used: Mouse monoclonal anti-GFAP (Sigma-Aldrich G3893 1:1000 dilution), Rabbit anti-Iba1 (Wako; 019-19741, 1:500 dilution); Rabbit anti-cleaved-Caspase-3 (D175) (Cell Signaling; 9661S 1:250 dilution); anti-HSP70/HSP72 (C92F3A-5) (Enzo; ADI-SPA-810-F 1:400 dilution). Slices were mounted in VECTASHIELD Antifade Mounting Medium with DAPI (Vector labs; H-1500). Imaging was performed with a Pannoramic 250 Flash slide scanner (3DHistech) at 10X magnification. Resulting images were analyzed with a custom Matlab script. Regions of interest ~1mm wide and encompassing the depth of the neocortex were selected at the center of the heating site and in the mirror position in the contralateral hemisphere. Mean fluorescence intensity on the treated side was divided by mean intensity on the contralateral side, for each label.

**Statistical analysis.** Statistical analyses were performed in Matlab using the `fitglme` and `fitrm` functions (Statistics and Machine Learning toolbox) to fit ANOVA models, and `ttest` for paired twotailed t-tests.

## Results

### Brain temperature under a craniotomy

We first measured factors affecting the baseline temperature of the mouse brain. Optical experiments typically involve replacing a section of skull with a glass coverslip to provide access to the brain. The removal of the overlaying bone, skin, hair, and vasculature, and implantation of a glass window and metal headpost affect thermal equilibrium at the brain surface.

We measured the brain surface temperature of freely-moving mice without a craniotomy, using a thermocouple probe (220 μm diameter) implanted through a small burr hole above the visual cortex. The temperature was 36.8°C 30 minutes after recovery from anaesthesia (N=2 mice), consistent with previous reports (Shirey et al., 2015).

We next measured brain temperature under a cranial window using a thermocouple probe inserted through a gap at one edge of the glass (Fig. 1a). Brain temperature 500 μm below the cranial window was 35.3°C in awake headfixed mice in the absence of objective immersion, which reduced to 34.3°C with room-temperature immersion water placed above the window and a microscope objective immersed (N=2 mice). Anaesthesia induces prolonged brain cooling for up to 30 minutes after recovery (Shirey et al., 2015). We found that 1.25%isoflurane anaesthesia reduced brain temperature 500 μm below the dura to 33.8°C and 34.4°C in the presence and absence of immersion water, respectively, while core temperature was maintained at 37°C (N=2 mice). We also performed measurements 100 μm below the dura (Table 1). The measured temperatures are consistent with the degree of cooling previously described at the brain surface (Kalmbach and Waters, 2012).

**Table 1.** Cranial temperatures measured with a thermocouple in the absence of laser illumination. In the *No window* condition, a thermocouple was implanted through a burr hole in the skull, with skin sutured over the site. In the other conditions, a window was implanted and headbar affixed to the skull Awake measurements were performed 30 minutes after recovery from anaesthesia. Mouse body temperature was maintained at 37°C. Depths relative to dural surface. Multiple values within a cell denote biological replicates. *nm*: not measuredFigure 1

| Condition |  | Depth | Temperature (°C) |  |  |
| --- | --- | --- | --- | --- | --- |
|  |  |  | Awake<br>Freely moving | Awake<br>Headfixed | Anaesthetized<br>(1.25% isoflurane) |
| No window |  | 100 µm | 36.8<br>36.8 | *nm* | 34.8<br>36.0 |
| Partial window | No immersion | 100 µm | *nm* | 33.6 | 32.8 |
|  |  | 500 µm |  | 35.6<br>35.0 | 34.8<br>34.0 |
|  | Water + objective | 100 µm | *nm* | 32.4 | 32.2 |
|  |  | 500 µm |  | 34.7<br>33.9 | 34.2<br>33.4 |
| Full window | No immersion | Glass surface | *nm* | 32.4 | 31.7 |
|  | Water + objective | Glass surface | *nm* | 29.8 | 29.5 |

### Brain heating during two-photon imaging

We performed contact temperature measurements of laser-induced heating using the thermocouple probe inserted under a partial window (Fig. 1a). In all experiments, we raster scanned (30 Hz) a 1mm square region 250 μm below the brain surface. We performed experiments with highly focused (NA=0.8) or weakly-focused laser light (NA=0.08, achieved by underfilling the objective) to produce intensities characteristic of point scanning or parallel illumination schemes, respectively. Upon laser illumination, the thermocouple registers a temperature change due to heating of the surrounding tissue as well as direct absorption of light by the thermocouple itself. To isolate heating in the tissue, we applied a heat transfer model consisting of four compartments, described by a set of ordinary differential equations with five free parameters (Fig. 1b, Methods). Fitting this model to thermocouple temperature measurements allows recovery of the temperature timecourse of the illuminated brain volume, compensating for the effect of the thermocouple (Fig. 1c). Here we report temperature changes of the illuminated brain volume, compensated for the effect of the thermocouple, after 20 seconds (ΔT_20_) and 180 seconds (ΔT_180_) of illumination. Twenty seconds corresponds approximately to the time constant of heating, whereas at 180 seconds temperatures approached steady state.

Numerical aperture had no effect on heating rates when scan area and focal plane depth were held constant (Fig. 1d,e). To reduce effects of nonlinear absorption and allow higher average powers to be tested, we performed the remaining experiments reported in this section with weakly-focused light.

We recorded temperature changes at illumination powers up to 400 mW at wavelengths of 800,920, and 1040 nm. Temperature changes were consistent with a linear proportional relationship between illumination power and heating (Fig. 1f,g). Longer wavelengths produced slightly greater heating (p<10^−5^, ANOVA), but heating varied much less than water absorption (Curcio and Petty, 1951) at these wavelengths, consistent with the majority of light being absorbed before escaping the brain. We measured the spatial spread of heating by keeping the thermocouple fixed and varying the location of the scanned area along the antero-posterior axis of the brain. The temperature change decreased with distance, but remained detectable (30%of maximum) at 4.8mm, the maximum distance we measured (Fig. 1i). The time constant of heating was prolonged at greater distances, resulting in a narrower spatial scale of heating 20 seconds after illumination onset (Fig. 1h) compared to 180 seconds (Fig. 1i).

We next performed independent, non-contact temperature measurements using fluorescence emission spectrometry of quantum dots. Quantum dots are brightly fluorescent semiconductor nanocrystals with temperature-dependent emission spectra (Li et al., 2007), which have been used for optical thermometry (Li et al., 2007; Maestro et al., 2011). Nanocrystals have previously been used to measure temperature under conditions inaccessible to contact thermometers, such as within living cells (Kucsko et al., 2013; Yang et al., 2011). We measured the dependence of steady-state brain temperature upon illumination power (ΔT_inf_) by measuring spectral shift across illumination powers of 30-450 mW (Fig. 2c–e) and comparing to spectral shifts obtained in calibration experiments *in vitro* (Fig. 2a). Quantum dot measurements in four mice indicated a mean ΔT_inf_ of 1.86±0.25 °C/100mW (mean±standard error) for 920nm illumination at a depth of 250 μm (Fig. 2e). This result was consistent with corresponding thermocouple measurements (ΔT_180_ = 1.50±0.02 and 1.73±0.04 at 150 μm and 500 μm respectively), noting that steady-state temperature changes (1000 seconds after onset) were 6%higher than those after 180 seconds. The temperature rise was similar whether mice were awake or anaesthetized (p=0.16, ANOVA; thermocouple measurements: N=3 mice; Qdot measurements: N=2 mice).

### Heating is greatest deep below the imaging plane

Due to scattering and absorption as light travels through tissue, total light power is highest at the brain surface. However, resting temperature is lowest at the brain surface due to the cranial window. Moreover, heat conduction through the coverslip reduces temperature changes at the brain surface, causing the greatest increases in temperature to occur deeper within tissue. To visualize spatial patterns of temperature and temperature changes in the brain we used a light-induced heating model to simulate imaging with focused light through an objective and cranial window (Fig. 3a–c) (Stujenske et al., 2015). This model incorporates anisotropic scattering and absorption of propagating illumination light, and simulates metabolic heating, circulation, and heat conduction using finite differencing methods. Simulations indicated that maximum levels of both absolute temperature (T, Fig. 3a) and heating (ΔT, Fig. 3c) occur approximately 500 μm below the brain surface, despite greater total absorption at the surface of the brain (Fig. 3b). To test these predictions, we performed temperature measurements with the thermocouple probe at different depths, observing temperature changes 20±7%higher (mean±std; Fig 3d–e) at 500 μm than at 150 μm. These results indicate that heating is more pronounced below the image plane than at the brain surface.

**Figure 3.**
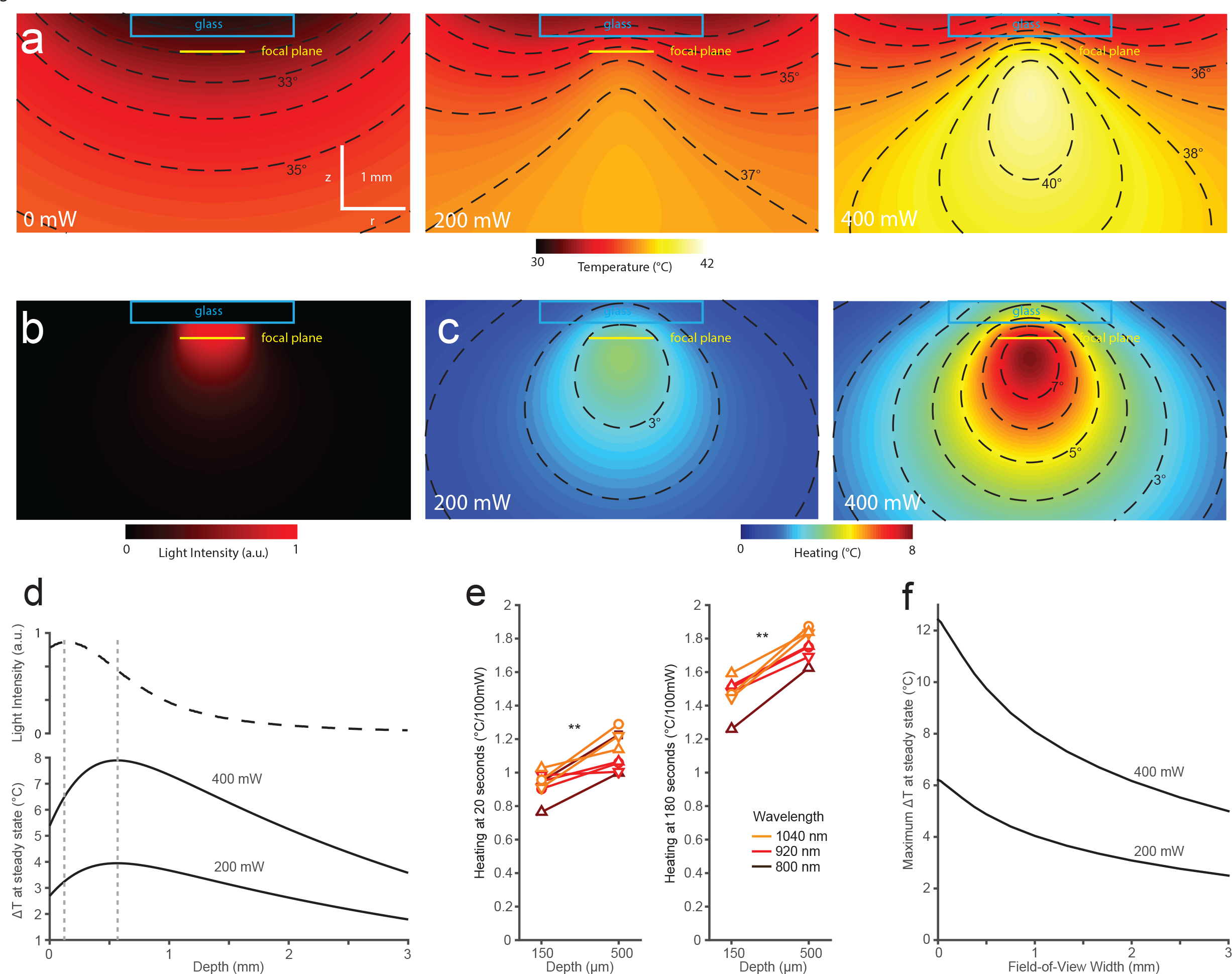
Heating is greatest deep below the brain surface. a) Simulated spatial temperature profile in brain tissue with overlaid cover glass and immersion water. The temperature of the immersion water was held at 25°C 2.5mm above the window glass. Profiles are shown for no illumination, 200 mW, and 400 mW of laser power raster scanned as in (b). b) Time-integrated light intensity cross-section through the center of the scan volume for the 1 mm square scan at 920 nm used in simulations. c) Simulated temperature changes at 200 and 400 mW. Contour lines are shown for 1°C intervals in (a-c). d) (top) Simulated time-integrated light intensity in a 150 um radius of the scan area center, at varying depth from the brain surface. (bottom) Steady-state heating in the same 150um radius, shown for 400mW and 200mW illumination powers. e) Heating coefficients measured by thermocouple contact thermometry at depths of 150 and 500 μm below the dura. (N=3 mice). f) Maximum temperature change for 200 mW simulated square scans of different field widths **:p<0.01, ANOVA

We used the same simulation to estimate how the size of the scanned area affects heating (Fig. 3e). At a given power, scanning a smaller area concentrates energy and produces a greater peak temperature, though this is offset by the rapid conduction of heat away from a smaller volume. Compared to the 1mm field of view, a simulated scan area 4 times smaller (0.5 mm) resulted in temperature changes 20.7%higher, while a 4x larger area (2 mm) resulted in 23.6%smaller changes. This relatively weak dependence of peak temperature on the illuminated area is consistent with the observed broad diffusion of heat within tissue.

### Tissue damage produced by laser heating

We assessed lasting heating-induced damage by scanning illumination at various power levels (0-450 mW) in awake mice. We extracted the brain 16 hours after imaging, and labelled fixed brain slices for signs of microglial activation (anti-Iba1 immunostaining), astrocytic activation (anti-GFAP immunostaining), heat shock (anti HSP-70/72 immunostaining), and activation of apoptotic pathways (anti activated Caspase-3 immunostaining). Immunoreactivity for Iba1, GFAP, and HSP70/72 increased strongly in the illuminated region following continuous illumination (20 min) at high powers. The spatial pattern of staining corresponded closely to that of temperature in our simulations (Fig. 4a, compare Fig. 3a). Increased immunoreactivity was evident in mice illuminated at 300 mW or above (Fig. 6a, Z-test, p<0.05). The mean intensity of activated Caspase-3 immunolabeling did not increase following heating, but punctate labeling of cells was observed at powers of 300 mW and above (Fig. 4a). No indications of persistent response to heating were seen at 250 mW or lower powers (Fig. 5, 6a). Control mice having a window implant but no laser heating showed little or no increase in immunoreactivity below the craniotomy (Fig. 6a, Iba1:1.08±0.03, GFAP: 1.20±0.10, HSP: 1.22±0.13, Caspase-3: 0.95±0.04, mean±standard error of fractions vs opposite hemisphere).

**Figure 4.**
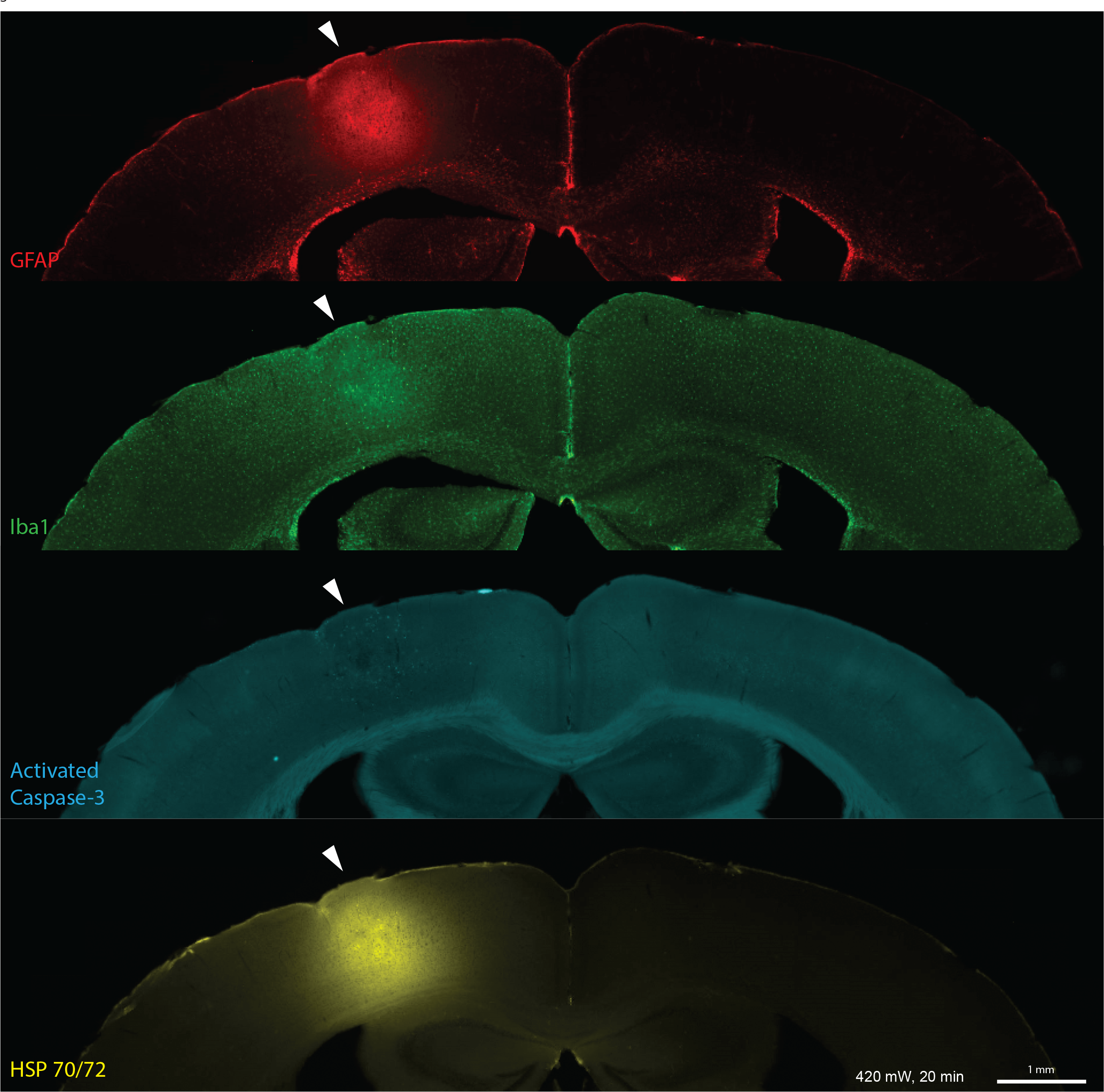
Immunolabeled sections of a mouse brain illuminated at 420 mW, 920nm, fixed 16 hours after heating. The location of heating is indicated with white arrowheads.

**Figure 5.**
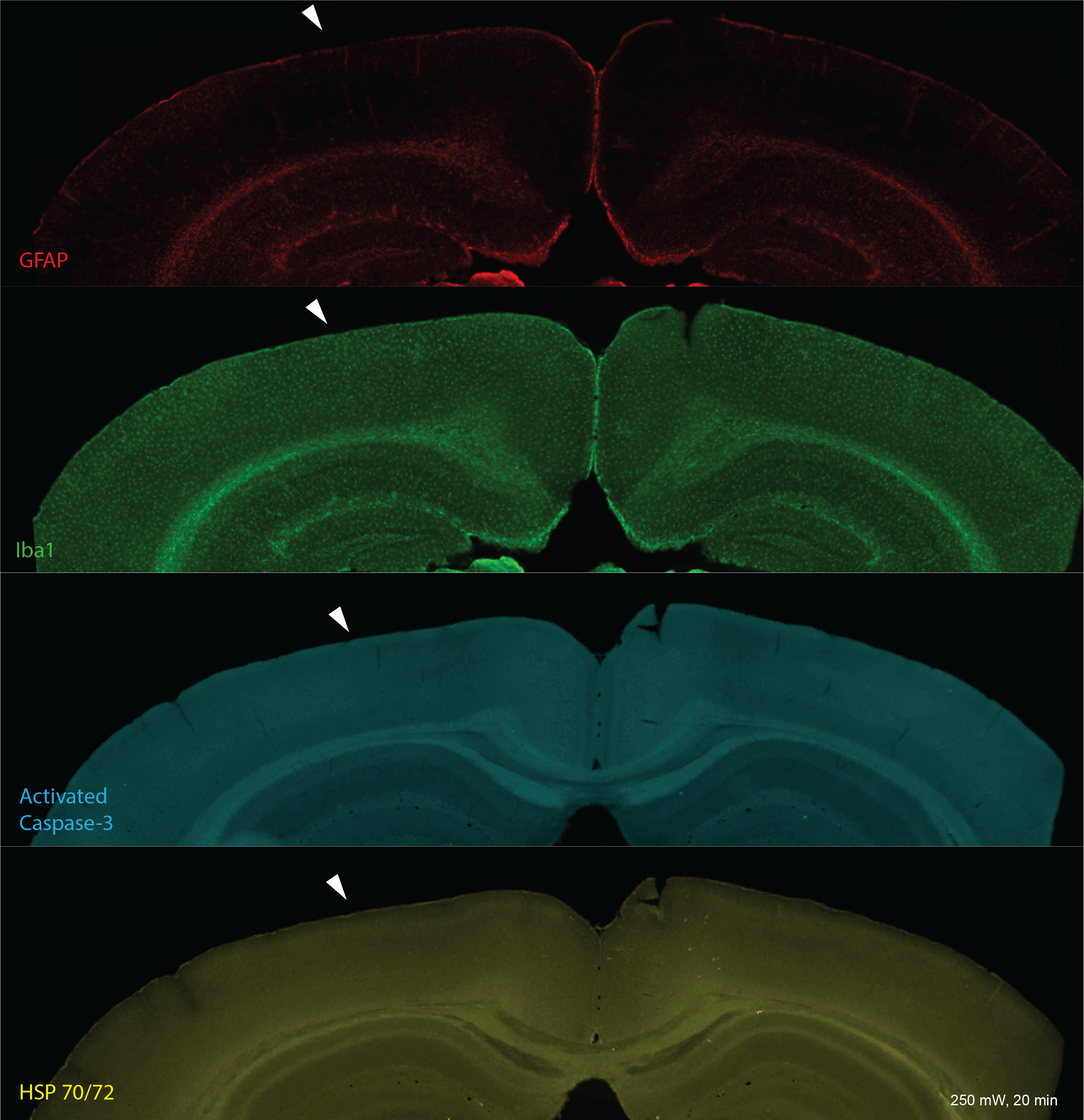
Immunolabeled sections of a mouse brain following 250 mW continuous illumination for 20 minutes, 920nm, fixed 16 hours after heating. The location of heating is indicated with white arrowheads.

Heating-induced damage is not linear in the input power, with small increases in temperature above an onset threshold causing rapid increases in damage, likened to the activation energy of other chemical processes (Dewey, 2009). We reasoned that reducing illumination duty cycle to periods much shorter than the time to reach steady state would reduce the peak temperature achieved, permitting higher optical powers with less heating damage. Using a 33%duty cycle (10 sec on, 20 sec off) for 1 hour, we observed no signs of histological damage at illumination powers below 400 mW (Fig. 6a). Immunoreactivity for GFAP, HSP, and Iba1 was significantly reduced at the lower duty cycle (MANCOVA, p<0.05).

**Figure 6.**
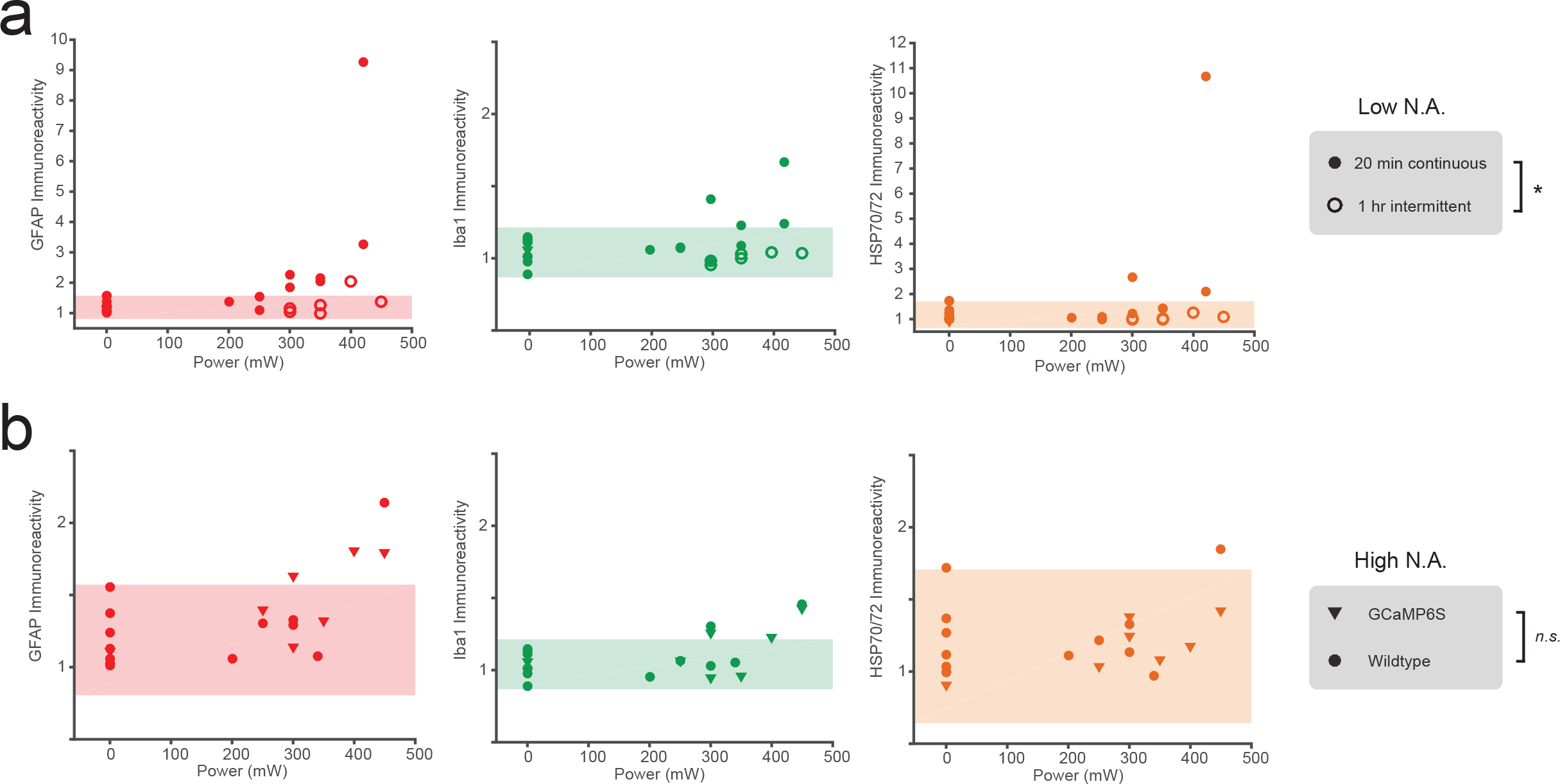
Quantification of heating-induced damage. a) Intensity of immunolabeling, as fraction of intensity in contralateral hemisphere, in mice heated at different laser intensities, under continuous (20 min; closed circles) or intermittent (1hr, 33%duty cycle, 10s pulses; open circles) illumination. The shaded area denotes the 95%confidence interval of the control group mean. b) Intensity of immunolabeling, as fraction of intensity in contralateral hemisphere, in GP4.12 (triangles) or Wild-Type (circles) mice heated at different laser intensities for 20 minutes. The shaded area denotes the 95%confidence interval of the control group mean. Control data (0 mW) are shared in a) and b). *: p<0.05, MANCOVA. ns: not significantHeating at 20 seconds (°C/100mW)

We also assessed damage when scanning a highly focused beam (NA=0.8). Because photodamage can be mediated by fluorophore excitation, we conducted these experiments with both wild-type mice and GP4.12 (GCaMP6S) transgenic mice (Dana et al., 2014). As with weakly focused light, increased immunoreactivity was evident at powers of 300mW and above in both mouse strains following 20 min continuous illumination (Fig. 6b). No signs of localized damage near the focal plane were observed for illumination powers below this threshold in either mouse strain. These results suggest that heating, and not nonlinear damage, determines the maximum usable power for the optical configuration used here.

## Discussion

Laser light used for two-photon imaging and optogenetics is absorbed by brain tissue, and can significantly raise brain temperatures at high intensities. For a 1mm^2^ scanned region, this heating corresponds to a maximum of approximately 1.8°C/100mW. Larger scan areas result in smaller temperature changes, and smaller areas result in larger changes. However, this dependence is relatively weak, because the spatial extent of heating is broad, spanning millimeters. Although most light is absorbed near the brain surface, peak heating occurs deeper in the brain, because the brain surface is cooled by conduction through the coverslip to the immersion fluid. For a focal depth of 250 μm, peak heating occurs at roughly 500 μm. We expect these results to apply to a wide range of scanning or illumination patterns due to the broad spread of heat within brain tissue.

The goal of these experiments was to establish practical limits to illumination power for two-photon excitation experiments. Heating is particularly relevant to experiments employing multiple focal points (e.g. Bewersdorf et al., 1998; Bahlmann et al., 2007; Amir et al., 2007; Cheng et al., 2011; Papagiakoumou et al., 2013; Voigt et al., 2015; Packer et al., 2015), in which high total power is a greater concern than peak power at any one focus. The amount of heating tolerable in a given experiment depends on many factors, such as the duration and duty cycle of the experiment, and the physiological variables being measured. For example, power limits may differ in anaesthetized versus awake animals due to the neuroprotective effects of anaesthesia (Sanders et al., 2005) and its concomitant reduction in brain temperature (Shirey et al., 2015; Yenari and Han, 2012). Here we investigated the threshold for lasting histological damage in awake mice, finding damage onset in the range of 250-300 mW for 20-minute continuous illumination, corresponding to a steady state temperature rise of approximately 5°C. Larger powers were tolerated at a lower duty ratio. This decrease in tissue damage at the reduced duty cycle indicates a threshold phenomenon, because the cumulative illumination dose was identical between the two protocols.

The damage thresholds measured here are upper bounds. Transient changes in physiology may occur at lower powers. For example, changes in neuronal firing rates are reported *in vivo* with illumination-induced heating of approximately 1°C (Stujenske et al., 2015). A possible choice of power limit is defined by natural variations in brain temperature induced by activity. Salient stimuli and physical activity are accompanied by transient increases in brain temperature, with changes of 3°C reported in rodents (Kiyatkin, 2005).

Recent technical advances have introduced simultaneous imaging in multiple brain regions (Lecoq et al., 2014; Voigt et al., 2015), raising the question of whether total power delivered to the brain can be increased if focal regions are sufficiently separated. Our results indicate that temperature changes spread within tissue over several millimeters, and similarly large spacings would be required to achieve more than modest increases in the total power deliverable to the brain.

Our experiments also inform the optimal degree of multiplexing in configurations that illuminate multiple focal spots in a single focal region. The signal produced by *n* foci, each with power *P*, is *nP*^2^ (omitting scaling). Given a maximum power deliverable to a single focus, *P_local_*, limited by nonlinear damage, and a maximum total power, *P_global_*, limited by heating, the signal rate *S(n)* achievable by *n* foci is determined by whichever threshold is met first: 
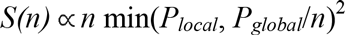
*S(n)* is maximized by selecting *n*=*P_global_*/*P_local_*, *i.e*. when the local and global damage thresholds are met simultaneously. If *n* were higher, more signal could be achieved by concentrating limited total power into fewer spots, increasing quadratic excitation. If *n* were lower, the total power used would be below the heating threshold, so more signal could be achieved by adding a focus. The optimal degree of multiplexing in parallel microscopy systems therefore depends on tradeoffs between total power (more focal points) and peak power (fewer focal points). The present study addresses only global power limitations, which are broadly applicable, while nonlinear damage depends strongly on the details of the optical configuration (Ji et al., 2008; Donnert et al., 2007; Kawano et al., 2003; Hopt and Neher, 2001; Koester et al., 1999).

The volume and time over which illumination is distributed is an important consideration for laser-induced damage. Focused ultrafast laser pulses cause heating over short length and time scales. However, because diffusion dissipates heat rapidly over short length scales, transient temperature changes in the focal volume produced by laser pulses over a typical pixel dwell time are small compared to the tissue heating described here over longer spatial and temporal scales (Schönle and Hell, 1998). In our low N.A. experiments, laser pulses were not delivered to a sharp focus, but distributed over an area in the focal plane approximately 100 times larger than the diffraction limit of the objective, which was also rapidly scanned in space (see Methods). Furthermore, the spatial profile of the damage closely overlapped measured patterns of heating, peaking well below the focal plane. It is therefore unlikely that the histological damage we observed at high powers was due to effects at the beam focus.

These experiments explored a limited space of surgical and laser scanning protocols. Heating and damage may differ in other cortical regions, other preparations (*e.g.* a thinned skull (Yang et al., 2010) or open craniotomy (Xu et al., 2012)), or where excitation is limited to very short spatial or temporal scales (e.g. single-neuron spike injection (Packer et al., 2015)). In other preparations, heating parameters may differ or other damage mechanisms may dominate. Although heating does not differ between highly and weakly focused beams, photodamage can be affected by the spatial pattern of illumination. Far from the focal plane, a weakly focused beam produces higher peak intensities than a highly focused beam, which could contribute to high immunoreactivity ratios we observed with weakly focused light. Under parallel illumination, each sample volume is illuminated for a larger fraction of time, which could recruit damage mechanisms differently than under highly focused light. Conversely, highly focused light can induce significant focal damage particularly when scanned over small areas (Hopt and Neher, 2001; Ji et al., 2008; Koester et al., 1999). Interactions between temperature and nonlinear effects can also occur (Chirico et al., 2003). As a result of the diverse factors affecting photodamage, these damage thresholds do not necessarily extend to other illumination conditions.

We demonstrated that damage can be reduced for a given peak power and total light dose by using lower duty ratios, which result in lower peak temperatures within brain tissue. An alternative approach to controlling brain temperature takes advantage of strong heat conduction through the cranial window (Kalmbach and Waters, 2012). Room temperature immersion results in significant brain surface cooling, and lowered temperatures are only compensated in the focal plane at roughly 200 mW power, likely mitigating negative effects of heating. Appropriate temperatures could be maintained for higher or lower powers by perfusion of temperature-controlled immersion fluid (Kalmbach and Waters, 2012), potentially enabling higher illumination powers without damage.

A related concern is that neurons adapt to chronic temperature changes by regulating membrane conductance (Magee and Schofield, 1991). Our experiments indicate that the chronic cortical surface temperature of mice in their home cage (*i.e.* in the absence of immersion) is reduced by 1.5°C below a cranial window implant. It remains unknown how the brain adapts to such changes, or how adaptation would interact with subsequent heating during imaging.

In conclusion, our results describe heating effects induced by two-photon imaging and their dependence on several illumination parameters, using two independent thermometry techniques and immunohistochemistry. High illumination powers are increasingly being used to apply optical methods on unprecedented spatial and temporal scales. We have identified damage thresholds for such experiments and demonstrated procedures that avoid harmful effects.

## Acknowledgements

The authors thank Amy Hu, Alexander Song, Hod Dana, Simon Peron, Kayvon Daie, Jared Rouchard, John Macklin, Mladen Barbic, Karel Svoboda, Na Ji, and Jeffrey Magee for assistance with these studies and comments on the manuscript. Laboratory space and equipment for these studies were provided by Karel Svoboda and Jeffrey Magee.

